# Functional profiling of the sequence stockpile: a review and assessment of *in silico* prediction tools

**DOI:** 10.1101/2023.07.12.548726

**Authors:** R Prabakaran, Y Bromberg

## Abstract

*In silico* functional annotation of proteins is crucial to narrowing the sequencing-accelerated gap in our understanding of protein activities. Numerous function annotation methods exist, and their ranks have been growing, particularly so with the recent deep learning-based developments. However, it is unclear if these tools are truly predictive. As we are not aware of any methods that can identify new terms in functional ontologies, we ask if they can, at least, identify molecular functions of new protein sequences that are non-homologous to or far-removed from known protein families.

Here, we explore the potential and limitations of the existing methods in predicting molecular functions of thousands of such orphan proteins. Lacking the ground truth functional annotations, we transformed the assessment of function prediction into evaluation of functional similarity of *orphan siblings*, i.e. pairs of proteins that likely share function, but that are unlike any of the currently functionally annotated sequences. Notably, our approach transcends the limitations of functional annotation vocabularies and provides a platform to compare different methods without the need for mapping terms across ontologies. We find that most existing methods are limited to identifying functional similarity of homologous sequences and are thus descriptive, rather than predictive of function. Curiously, despite their seemingly unlimited by-homology scope, novel deep learning methods also remain far from capturing functional signal encoded in protein sequence. We believe that our work will inspire the development of a new generation of methods that push our knowledge boundaries and promote exploration and discovery in the molecular function domain.

## Introduction

A typical cell contains about 0.2 g/ml proteins, which translates to up to a billion molecules per cell^1, 2^. However, the corresponding number of distinct protein sequences varies from only a few hundred in some bacteria to tens of thousands in many eukaryotes. Characterizing these vital biomolecular nanomachines, i.e. identifying their cellular functions, associated pathways, localization, interaction partners, and catalytic activities, is crucial for understanding their role in cellular biology. Experimental annotation of protein function remains significantly limited by its cost and speed. For example, among the 94.5 million protein sequences that have been deposited in UniProt in the last three years, only 6,974 (<0.01%) were manually curated. Thus, the growing influx of sequencing data has necessitated accurate computational annotation of protein function for diverse downstream analyses.

Over the last two decades, the number of bioinformatics tools developed for *in silico* protein annotation has grown and algorithms diversified. Historically, the most common and reliable techniques for annotation relied on the transfer of function by homology, i.e. shared ancestry resulting in sequence similarity. To characterize a given query protein, various alignment and domain profiling tools such as BLAST, PSI-BLAST, and HMMER^3–7^ were used to search annotated protein databases^8–11^. More recently, faster algorithms have been developed to process and annotate large sequence datasets, including sequence reads and genes/proteins extracted from (meta)genome assemblies^12–16^. The challenges associated with protein functional annotation are multi-fold and have been discussed at length in earlier studies^17–20^. To summarize the state of the art: aside from defining what exactly the word “function” means in reference to proteins, there are three bottlenecks in producing accurate annotations – evolutionary caveats that limit function transfer by homology, lack of existing experimental annotations, and limitations of functional ontologies.

The first bottleneck arises as life evolves and adapts and divergent evolutionary processes result in homologous genes of different functions. These could end up as false positive functional annotations of sequence-and structurally-similar proteins^21^. One such example among many is the enzymatically inactive duck δ crystallin I that shares >90% sequence identity with the active δ crystallin II^21, 22^. At the same time, different genes converging to perform the putatively same function may have minimal homology – a false negative^23, 24^. For example, human (PDB:1PL8) and *Rhodobacter sphaeroides* (PDB:1K2W) sorbitol dehydrogenases are sequence different. Of course, we note that whether the human sorbitol dehydrogenase is functionally the same as its bacterial version is up for discussion. In general, diverged genes found in different species, i.e. orthologs, that do participate in the same molecular mechanisms, may not operate at the same rate or efficiency given the specific species’ environmental constraints – a functional difference that is often ignored. We argue that context in which the function is carried out should be thought of as part of the definition of function. However, this discussion is beyond the scope of this manuscript.

Second, by definition, the general dearth of experimental annotations is limiting for function transfer by homology. Furthermore, existing annotations are biased towards proteins from large families and to species of interest. For example, experimental evidence for GO annotations only exists for less than 15% of proteins in SwissProt^25^. The effects of these biases are compounded by the computational annotation of newly accumulated genomic data – a process that fosters annotation error propagation. Note that the existing functional annotations can, by default, only cover the observed part of the protein universe, i.e. annotation of new sequences may be flawed simply by our limited knowledge of biotic functional capacity (**Figure 1**). In short, the classical approach of transferring protein function by homology is complicated by convergent/divergent evolution, lack of experimental annotations, and errors in available computational annotations; it is also limited to existing classes of proteins, reducing chances of discovery of novel functions.

**Fig. 1:**
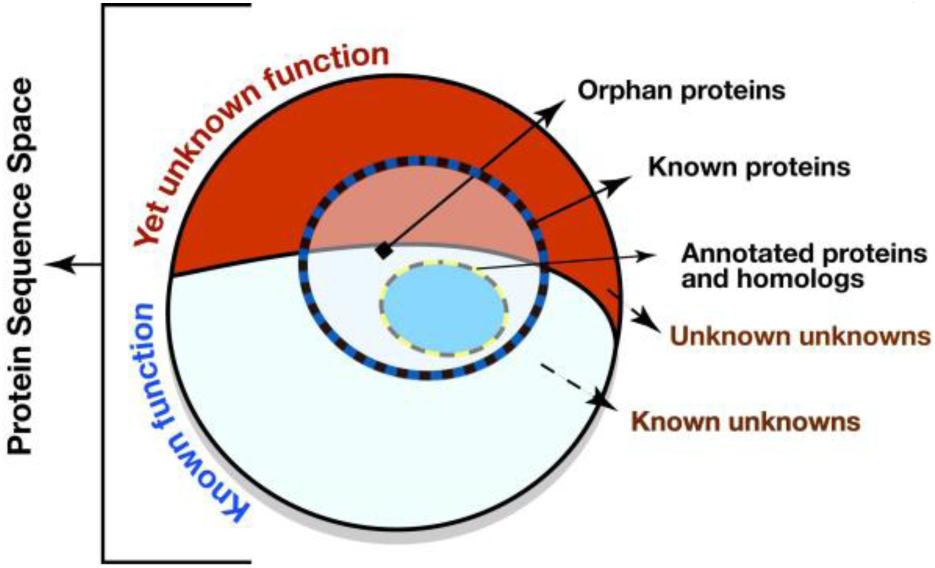
The limits of protein function annotations. Of the complete set of proteins (entire circle), containing known/previously observed proteins (blue/black dashed circle outline) and unknown/not-yet-seen proteins, some fraction carries out unknown functions (fraction of circle in red) rather than known (fraction in white) ones. Existing experimental and homology-based protein function annotations (blue oval) cover a small part of the complete protein sequence space. The number of orphan proteins, i.e. those lacking annotation and having no known homologs, is growing as we explore our world with better and faster gene/protein capture tools. Note that circle sizes are not to scale.

The third bottleneck is more technical in nature. The task of representing the ambiguous, environment-dependent, hierarchical role of a given protein with a set of human-understandable ontology terms is exceedingly difficult^11, 26–32^. Depending on the level of granularity and environmental conditions, a protein’s function could vary widely. For example, all kinases are phosphotransferases that catalyze the transfer of phosphate from ATP to carbohydrates, lipids, or proteins. However, kinases are part of almost every cellular process and many metabolic pathways, i.e. they can be assigned a wide range of biological functions. On the other hand, proteins involved in the same biological pathway have different catalytic (molecular) functions almost by definition; e.g. glycolysis (map00010;^27, 33^) involves kinases and dehydrogenases. That is, different molecular functions can contribute to the same biological role, while proteins of the same molecular function may have different biological roles – all across numerous environments and cellular compartments. An ideal protein function ontology should be robust to this variability, but also precise, widely applicable, expandable, and, lately, machine-readable. This ontology does not yet exist.

A significant amount of research has gone into targeting these challenges to computational function prediction. For examples, Critical Assessment of Functional Annotation algorithms (CAFA) is a community experiment that provides an even ground for the assessment of existing methods^34, 35^. CAFA employs a time-delayed evaluation where predictions of functions of a large set of yet-to-be-annotated genes/proteins are collected and assessed over a period of time through wet-lab experiments. CAFA results have documented the continuous emergence of new, better-performing prediction methods. Research has moved beyond sequence comparison, introducing new computational techniques, and incorporating additional biological data such as the protein-protein interactions, expression, phenotypic changes due to mutation, etc.

A key recent methodological development has been the ability to represent protein sequences as embeddings, i.e. projections of proteins into the latent space. Embeddings are interpretations of deep neural networks, learnt in the process of addressing a predefined objective function^36, 37^. Protein sequence embeddings have been successful in annotating various protein features, but most obviously protein structure^38–40^. Recently, deep learning methods have been developed to annotate protein function. For example, Littmann et al. have explored the application of protein embeddings to function annotation, reporting performance on par with CAFA ‘s top 10 best performers^41, 42^. Note that besides these representations capturing protein structural aspects and thus informing function, it remains unclear exactly which (or whether) aspects of functionality are reported by embeddings.

One important inference from the CAFA experience is the challenge of establishing metrics for the assessment of methods. That is, what is to be considered a correct annotation for a given protein? Given a protein P_1_ that carries out functions f_1_, f_2_, and f_3_ defined by a relevant ontology, if a method M_1_ predicts the protein to be of function f_1_ only, is this a correct annotation? How does this method perform in comparison to M2, which predicts the protein to carry out f_3_, f_4_, and f_5_? While for an individual annotation, say f_1_ *vs.* f_2_, ontology distance metrics can be established^43–46^, evaluating multiple annotations per protein is harder. Adding to the problem is the incomplete “ground truth” annotation, i.e. how would one take into account the protein’s unknown molecular functions?

Here, we provide a method and ontology-blind assessment approach for comparison of function annotation tools. We evaluate the predictions of computational methods for a set of proteins, sharing little sequence similarity with proteins in available databases. We ask, what is a correct annotation for a protein with no known sequence-similar homologs (i.e. an orphan)? To answer this question, we use structural similarity and a deep learning-based technique to establish whether a protein pair in our set shares functionality (i.e. are they siblings?), regardless of what specifically each protein does. We then evaluate other methods’ ability to recall shared functions for these pairs.

## METHODS

### Extracting the test dataset

From the ESM Metagenomic Atlas^47^, i.e. proteins translated from metagenome records of the MGnify database^48^, we collected 53,501,759 protein sequences, translated from metagenome-assembled genes, and having high-confidence predicted 3D structures, i.e. predicted local distance difference test (pLDDT) and predicted TM-score (pTM) greater than 0.9^38, 49, 50^. Note that our selected sequences make up less than a tenth of all structures in ESM and represent a significantly smaller fraction still of all metagenome-derived proteins MGnify collected over the years. Thus, the evaluation reported here is limited to a subset of ordered proteins, whose structure is well predicted and, thus, likely biased to reflect that of available, experimentally studied proteins.

These 53.5M sequences were aligned against UniRef100^51^ (using mmseqs2^13^ at default sensitivity =5.7). Sequences in UniRef are often used as reference for function transfer and as the training set for prediction models^9, 51^. To avoid using training data in our method testing, we focused on *orphan* sequences, i.e. those that do not share homology with proteins in UniRef. We identified 54,359 orphans with less than 30% sequence identity to UniRef100.

To further simplify the evaluation task, we filtered out longer proteins (over 400 residues) that are likely to contain multiple domains, as well as proteins whose sequences are truncated in the corresponding MGnify contigs, to retain 11,484 sequences. To avoid excess focus on sequence similarity, we further sequence-reduced this set at 90% identity using CD-HIT^52^. The final dataset of orphan proteins contained 11,444 proteins with ESM predicted structures and corresponding MGnify cDNA sequences.

### SNN + TM: annotating test set protein pairs as functionally similar (siblings)

In our earlier work^53^, we used structural similarity (TM-score^54^≥0.7) and functional similarity (SNN-score≥0.98, https://bitbucket.org/bromberglab/fusion-snn/), i.e. our SNN+TM approach (**Figure 2a**), to identify functionally identical enzyme pairs with 90% precision vs. the experimentally-determined Enzyme Commission^32^(EC) number annotations. SNN is a Siamese Neural Network, trained to identify functionally similar genes irrespective of sequence similarity^53^. The SNN architecture consist of a) pretrained embedding layer from LookingGlass^55^, b) a LSTM layer and c) computation of distance between embeddings (**Figure S1**). TM-scores for protein structure alignments were computed using Foldseek^54^.

**Fig. 2:**
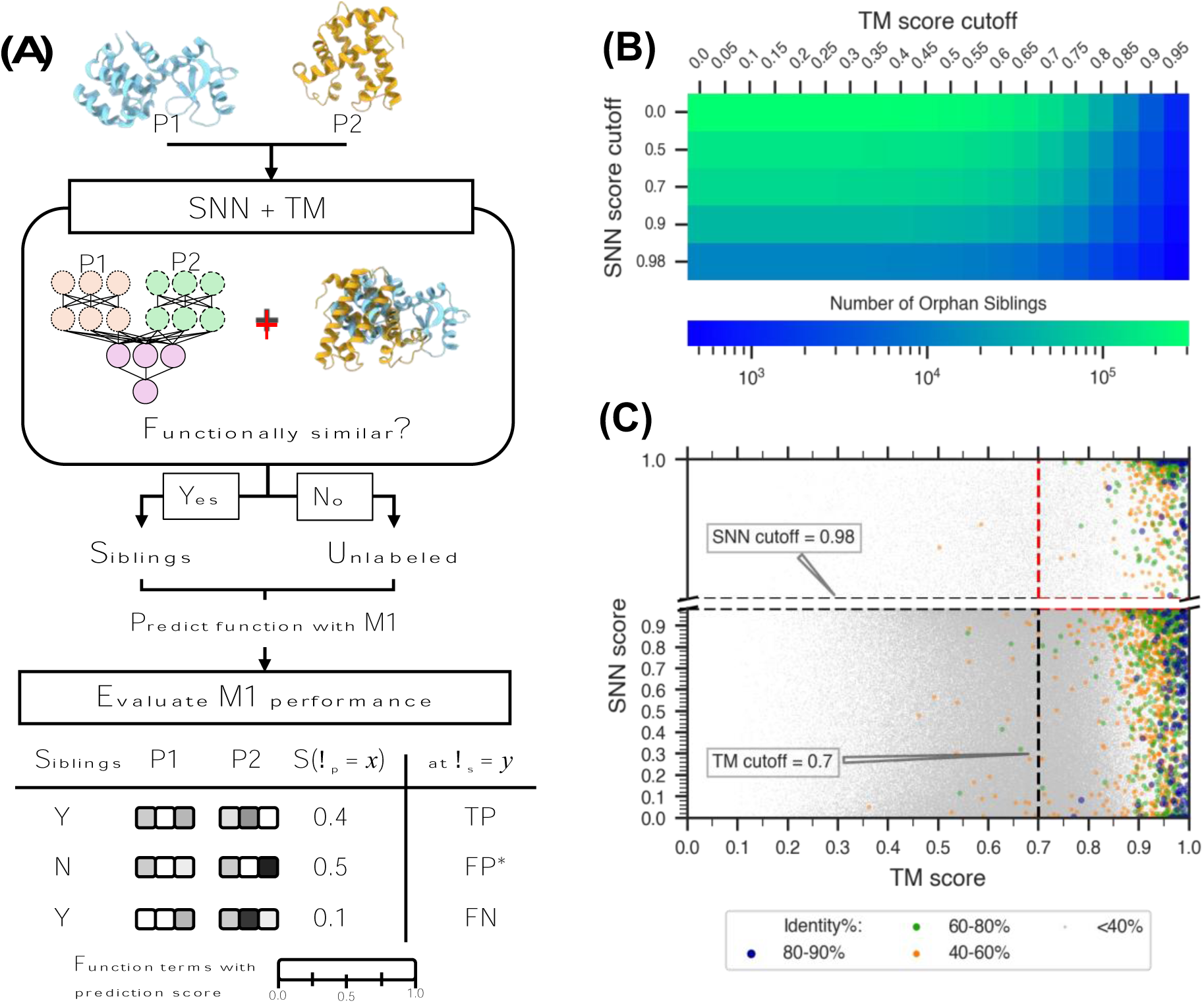
Evaluating function prediction through functional similarity. The performances of function prediction methods are evaluated based on the ability to predict functional similarity between protein pairs. (A) Putative functionally similar orphan protein pairs, i.e. a test set of orphan siblings, are built using the SNN+TM method. This SNN+TM test set of orphan siblings consists of protein pairs precisely labelled as siblings (pairs with high structure similarity [TM-score ≥ 0.7] and functional similarity [SNN score ≥ 0.98]) among unlabeled protein pairs. The performance of function prediction methods was evaluated by computing true positives, false negatives, and putative false positives (Methods). All assessments were conducted by varying thresholds for annotation prediction score (τp) and annotation set similarity (τs). (B) The number of orphan pairs considered as siblings at each threshold of TM score (x-axis) and SNN score (y-axis) on the plot is highlighted according to a log-scale gradient scheme from few proteins (blue) to many proteins (green). In (C) each dot represents a protein pair and is colored by sequence identity from very low (< 40%; gray) to very high (80-90%; dark blue). Note that no pairs over 90% identity were included in our set. The dashed lines indicate the TM and SNN score cut-offs (0.7 and 0.98) chosen for this work.

Note that in our original SNN+TM evaluation we used the available protein PDB structures^53^. Here, we planned to use ESMFold^39^-predicted structures of the orphan proteins instead. To evaluate the validity of using predicted structures, we identified *siblings* among a set of 1,869 enzymes with experimentally defined EC numbers and high-confidence ESM predicted structures (pLDDT and pTM greater than 0.9). Trivially, due to the slow pace of deposition of experimentally curated sequences into repositories, this enzyme dataset significantly overlaps with the set of proteins used for the original evaluation of SNN+TM method^53^. To build this enzyme set, we extracted from UniProt^9^ 5,697 enzymes of length ≤400 residues, annotated with a single, experimentally-evidenced EC number. Of these, only 33% (1,869) had high confidence ESMFold predicted structures.

We compared sibling annotations to EC pairings. The identified sibling pair was deemed correct if the corresponding EC annotations (at third level) were the same. We found that using structure predictions instead of experimental data did not significantly reduce our SNN+TM method’s ability to identify proteins of the same enzymatic functionality (precision = 87.8% here vs ∼90% in the original estimate). Note that we also used this dataset to estimate the performance of an ideal function predictor.

We further extracted functionally similar pairs, *orphan siblings*, from our set of 11.4K orphan proteins. We ran Foldseek (with TM threshold=0) to compare the predicted structures of all 11.4K proteins in our set amongst themselves. Only 309,549 of these protein pairs (0.5% of ∼65M possible ones) were structurally similar enough for a complete alignment. We then annotated functional similarity (SNN) scores for these 309K pairs (**Figure 2**).

Only 6K (6,219, 2% of 309K) pairs attained the pre-set cutoffs (TM score≥0.7, SNN score≥0.98) for shared function and were thus labelled *orphan siblings*. Note that at these cutoffs, proteins that are not identified as being of the same function can still be functionally identical (recall = 2.4%). Thus, we primarily assessed the performance of function annotation tools based on their capacity to find all test set orphan siblings, rather than on their precision of labelling pairs as functionally similar. Note that as our method for identifying shared function is subject to selected thresholds, we also explored the performance of the selected prediction methods by varying the TM score and SNN score cut-offs (**Figures 4, S5** and ***SI Table 5***).

**Fig. 3:**
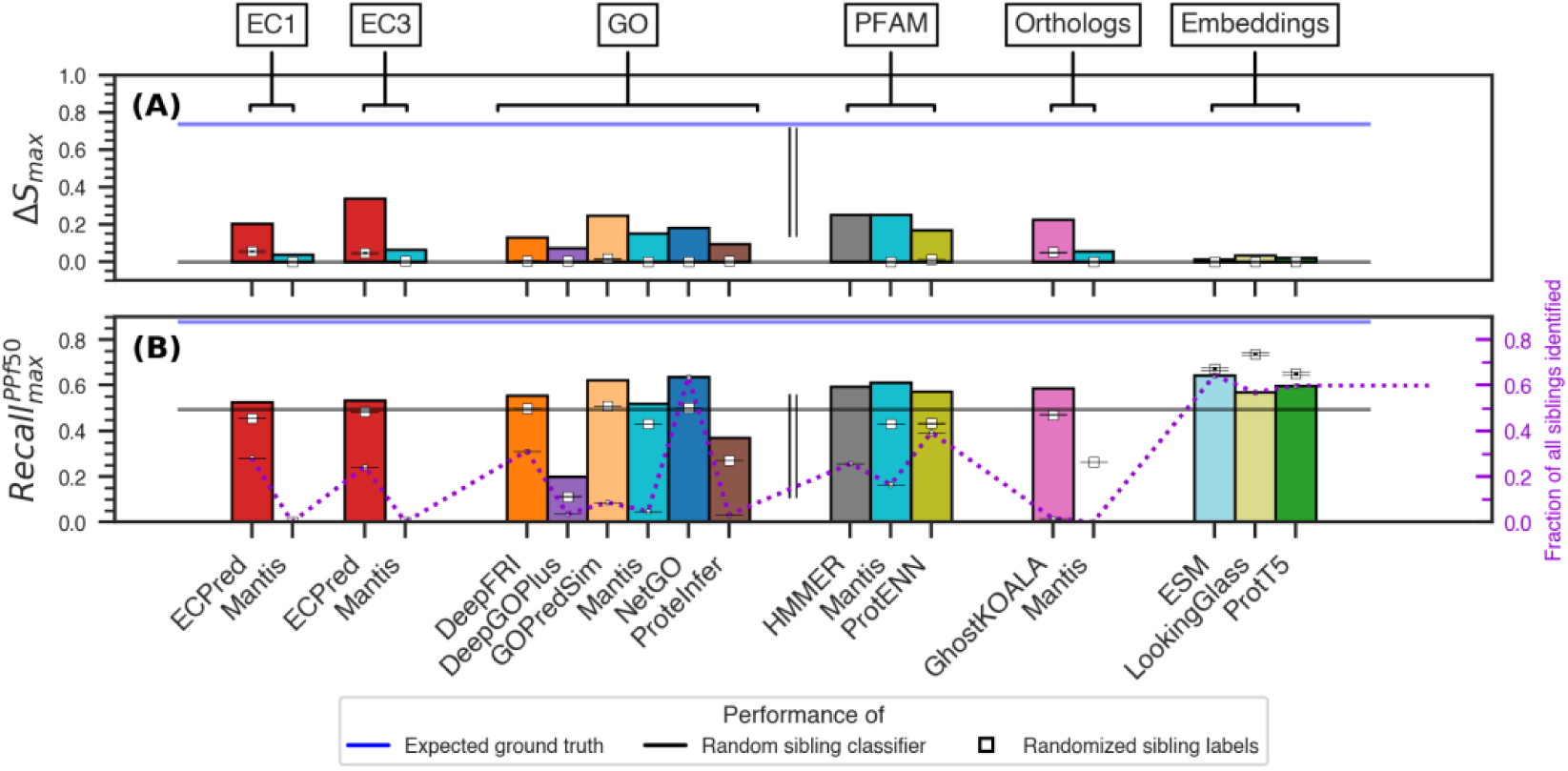
Comparison of prediction performance in each annotation category. The bar plots show the highest (max) per-method (A) ΔS and (B) **Recall^*ppf*50^** metrics for the SNN+TM test set of protein pairs (see **Methods**). The evaluated methods in this study predict: Enzyme Commission numbers (EC1 – first digit, EC3 – up to third digit), Gene Ontology Molecular Function terms, Pfam domains, and Ortholog groups (as indicated in (A). Language model embedding distances were also considered. Max scores were selected for each method by varying the prediction score threshold (τ_p_). The double line separates explicit functional annotations (EC and GO) from implicit functional annotations via Pfam and Ortholog definitions and embedding similarities. The gray and blue lines indicate the average performance of 100 random baseline classifiers and the expected performance of the “ground-truth” annotation (**Table S1**), respectively. The white squares along standard error bars indicate the average performance of the method-specific random annotators over 100 iterations.

**Fig. 4:**
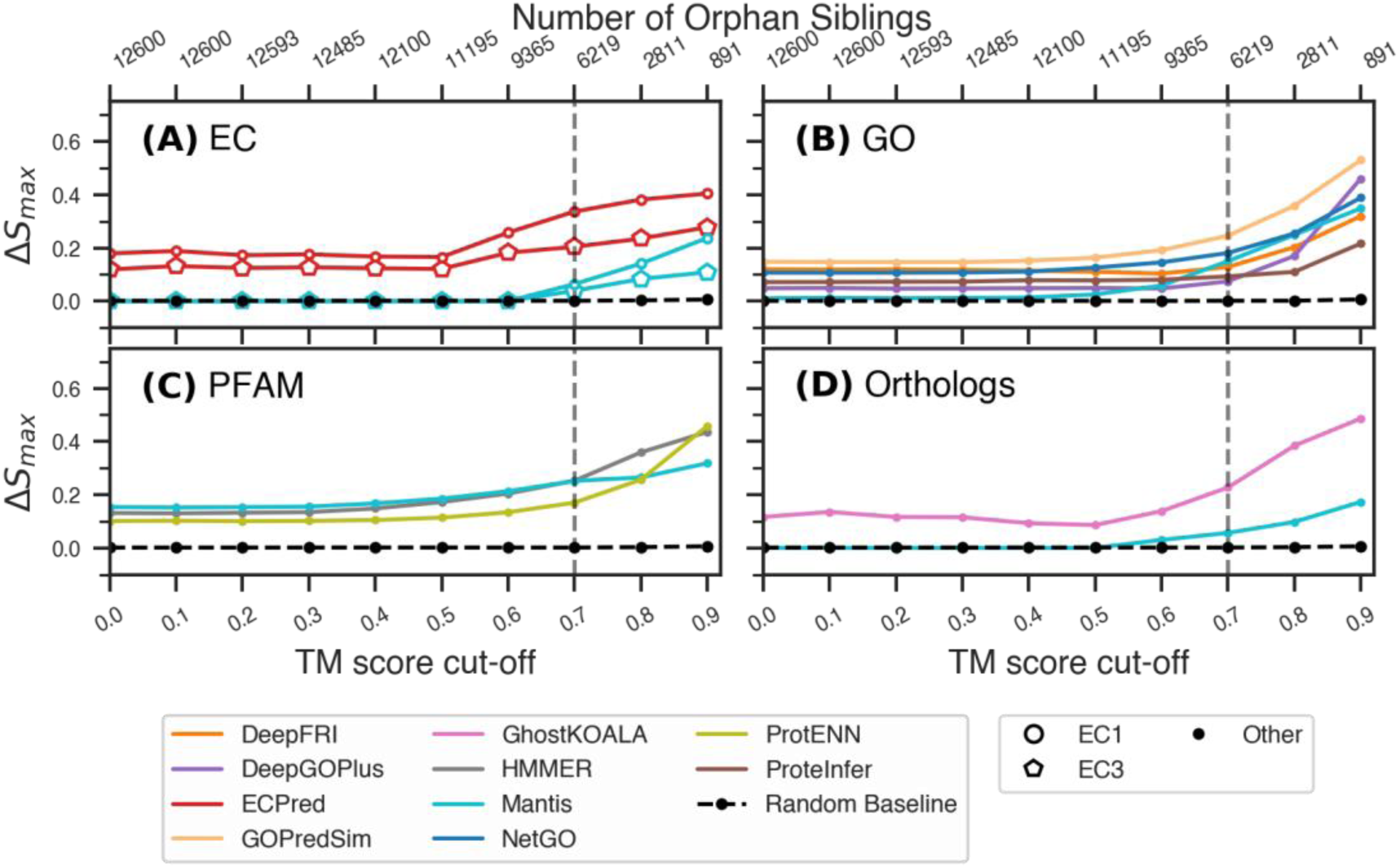
Variations in performance over TM-score cut-offs. The highest (A-D) ΔSimilarity (y-axis) of top performing methods vary based on predicted protein annotations. All scores were computed at SNN cutoff =0.98 and at different TM-score cutoffs (bottom x-axis), resulting in a different number of orphan sibling pairs (top x-axis). The average performance of 100 random baseline classifiers is plotted for comparison in each panel (dotted black line). See Figure S3 for trends of F1max and AUC PRmax.

### Translating predicted annotations into functional similarity

We reformulated the molecular function prediction challenge to overcome the limitations of evaluating and comparing methods that target different functional vocabularies, i.e. tools predicting Gene Ontology (GO) molecular function (MF) terms or Enzyme Commission (EC) numbers. We also included tools that identify protein Pfam domains and assign sequences to KO and COG ortholog groups^11, 56, 57^. Note that while GO and EC aim to explicitly describe the function of the protein, Pfam and Ortholog methods capture protein families and evolutionary relationships, which are related to, but not necessary directly reflective of function. If a protein was annotated with multiple Pfam domains, EC numbers, or orthologous groups, we retained all labels. We selected 13 protein annotation tools for our assessment based on the availability of a standalone version or a web server that can process multiple sequences. These 13 methods were grouped based on the type of protein annotation into four categories: GO – DeepFri^58^, DeepGOPlus^59^, GoPredSim^42^, GOProFormer^60^, and NetGO^61^, Pfam – HMMER^62^, InterProScan^63^, ProtENN & ProtCNN^64^ and ProteInfer^65^, Orthologs – GhostKOALA^66^, KofamKOALA^67^ and E.C. – ECPred^68^ and Mantis^69^. Methods such as Mantis^6, 69^ and ProteInfer^65^ were included in more than one category since they provide multiple types of protein annotations (**Table 1**). We also evaluated using pairwise DNA/Protein embedding similarities from unsupervised models such as Bepler^70^, CPCProt^71^, ESM-2^39^, LookingGlass Encoder^55^, ProtTrans^37^, SeqVec^72^ and Word2Vec^73^. In addition, we used SwiftOrtho^74^, a method that identifies orthologous pairs in a given set of proteins^75, 76^, as an upper bound of homology-based evaluation of our test set.

**Table 1:** Protein function prediction performance.

| Tool | Term | # of Annotated proteins<sup>1</sup> | % of annotatable protein pairs<sup>1</sup> | % of annotatable sibling pair<sup>1</sup> | $\tau_p^{\text{opt}}$ at $F1_{\text{max}}^2$ | $\tau_s^{\text{opt}}$ at $F1_{\text{max}}^2$ | $\Delta S$ | $F1_{\text{max}}$ | PR AUC | ROC AUC | Prec | Rec |
| --- | --- | --- | --- | --- | --- | --- | --- | --- | --- | --- | --- | --- |
| ECPred | EC1 | 9,402 | 60.1 | 73.3 | **0.800** | **0.01** | **0.18** | **0.67** | **0.72** | **0.60** | **0.57** | **0.87** |
| ECPred | EC3 | 9,402 | 60.1 | 73.3 | **0.890** | **0.01** | **0.25** | **0.68** | **0.70** | **0.67** | **0.65** | **0.75** |
| Mantis | EC1 | 1,910 | 8.5 | 6.3 | - | 0.19 | 0.04 | 0.64 | 0.71 | 0.52 | 0.51 | 0.90 |
| Mantis | EC3 | 1,910 | 8.5 | 6.3 | - | 0.19 | 0.05 | 0.61 | 0.65 | 0.53 | 0.52 | 0.76 |
| DeepFRI | GO | 11,328 | 97.1 | 98.3 | 0.214 | 0.01 | 0.06 | 0.47 | 0.59 | 0.60 | 0.62 | 0.44 |
| DeepGOPlus | GO | 11,444 | 100.0 | 100.0 | 0.100 | 0.02 | 0.01 | 0.23 | 0.40 | 0.55 | 0.65 | 0.15 |
| GOPredSim | GO | 11,444 | 100.0 | 100.0 | **0.384** | **0.05** | **0.14** | **0.63** | **0.66** | **0.61** | **0.57** | **0.72** |
| Mantis | GO | 2,895 | 11.8 | 12.6 | - | 0.38 | 0.10 | 0.59 | 0.60 | 0.58 | 0.56 | 0.64 |
| NetGO | GO | 11,444 | 100.0 | 100.0 | 0.087 | 0.48 | 0.07 | 0.53 | 0.67 | 0.68 | 0.62 | 0.66 |
| ProteinInfer | GO | 9,495 | 81.3 | 71.6 | - | 0.04 | 0.02 | 0.33 | 0.44 | 0.56 | 0.63 | 0.25 |
| GhostKOALA | KO | 3,551 | 7.3 | 9.7 | **0.696** | **0.01** | **0.11** | **0.65** | **0.70** | **0.57** | **0.55** | **0.85** |
| Mantis | COG | 5,784 | 59.9 | 47.2 | - | 0.10 | 0.04 | 0.60 | 0.64 | 0.53 | 0.52 | 0.75 |
| HMMER | Pfam | 8,661 | 83.5 | 76.3 | **4.400** | **0.64** | **0.22** | **0.69** | **0.76** | **0.66** | **0.58** | **0.89** |
| Mantis | Pfam | 4,874 | 15.0 | 26.9 | - | 0.37 | 0.16 | 0.62 | 0.65 | 0.64 | 0.62 | 0.65 |
| ProtCNN | Pfam | 11,444 | 100.0 | 100.0 | - | 0.01 | 0.00 | 0.13 | 0.42 | 0.52 | 0.77 | 0.08 |
| ESM | Emb | 11,444 | 100.0 | 100.0 | - | 0.05 | 0.00 | 0.19 | 0.72 | 0.70 | 0.25 | 0.07 |
| LookingGlass | Emb | 11,444 | 100.0 | 100.0 | - | 0.18 | 0.01 | 0.35 | 0.63 | 0.60 | 0.62 | 0.38 |
| ProtT5 | Emb | 11,444 | 100.0 | 100.0 | - | 0.08 | 0.00 | 0.22 | 0.72 | 0.71 | 0.35 | 0.12 |

<sup>1</sup>Unless explicitly specified, all our performance metrics of each method were computed within the respective subsets of the SNN+TM test set.
<sup>2</sup>Performance measures at $F1$ optimal thresholds are computed over 100 iterations of under sampling. Performance of the selected, top-performing methods is identified in bold. Precision and Recall were computed at the $F1$ optimum prediction score threshold ( $\tau_p^{\text{opt}}$ ) and optimum similarity score threshold ( $\tau_s^{\text{opt}}$ ). Other performance metrics: $\Delta S$ , $F1_{\text{max}}$ , PR AUC and ROC AUC are independent of the similarity score threshold ( $\tau_s$ ) by definition and thus were calculated at the $F1$ optimum prediction score threshold ( $\tau_p^{\text{opt}}$ ).

We scored the similarity between predicted annotations of proteins in each pair. Consider a protein pair P_1_ and P_2_ predicted by method M_1_ to carry out sets of functions Fu_1_ and Fu_2_, respectively. Fu_1_ (and Fu_2_) consist of several annotation terms from GO, EC, Pfam, or ortholog groups as assigned by M_1_; each term is associated with a prediction score (or E-value) and is accepted or rejected at a prediction score threshold (τ_p_). Increasing τ_p_ increases the precision of predicted annotations (Fu_1_ and Fu_2_) but could also reduce the number of predictions. All performance values reported in this study were computed by varying τ_p_ for methods that provide such prediction scores (**Table 1**). Note that, at a selected τ_p,_ different methods were only able to make predictions for subsets of our SNN+TM test set. Thus, all values reported here, unless explicitly specified, were computed on different sets of protein pairs.

The similarity (S) between Fu_1_ and Fu_2_ is defined by the Jaccard similarity coefficient, i.e. the ratio of the intersection set of terms to the union set (**Eqn. 1**). In case of GO annotations, we used the information accretion term^43, 77^ (I_a,_ **Eqn. 2,3**) to weigh the GO term assignment according to term frequency of appearance among UniProt GO annotations with experimental evidence^78, 79^; information accretion of a child GO term υ, I_a_(υ), is the information gained by adding υ to its parent term(s).

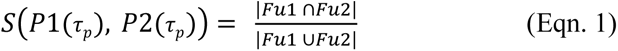

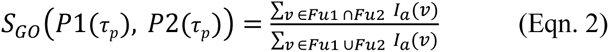

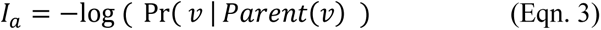

Additionally, the similarity score for a protein pair (P_1_ and P_2_) given their embeddings (E_1_ and E_2_) of length l, was derived from the Euclidean and cosine distances (**Eqn. 4 and 5**).

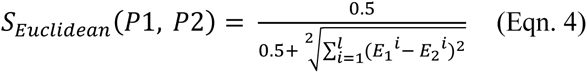

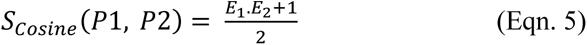

The performance of a given method in identifying *orphan siblings* was measured first by computing the Area under the Precision-Recall curve (PR AUC), area under the Receiver Operating Characteristic curve (ROC-AUC), and F1_max_ by varying the method-specific (**Eqn. 1, 2, 4 & 5**) similarity score threshold (τ_s_) for calling a protein pair functionally identical. To balance the number of sibling (positives) vs. unlabeled (mostly non-sibling) pairs, we under-sampled the latter to match the number of sibling pairs; we repeated the under-sampling 100 times and computed the average and standard deviation of all measures.

The F_1_ score, as the harmonic mean of Recall and Precision, provides an overall prediction performance in identifying siblings (**Eqn. 6**). We chose the optimum method prediction score threshold (τ^opt^p) and an optimum similarity score threshold (τ^opt^s) corresponding to the maximum F1 (F1_max_); we then reported Precision and Recall at these thresholds (**Eqn. 7-8**) for all methods. Note that for methods that predict EC numbers, PFAM domains and ortholog annotations, two proteins were considered to be functionally similar if they shared even one common annotation at a given prediction threshold (τ_p_).

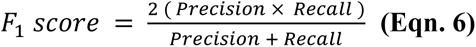

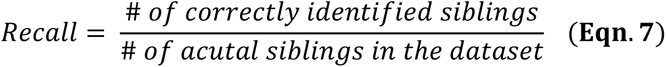

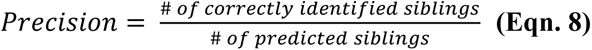

As described above, our SNN+TM approach has high precision, but a very low recall of functionally similar protein pairs. To compensate for the limitations of thus created test set, we report two additional performance measures: (1) the difference in similarity scores (ΔS, **Eqn. 9**) and (2) the maximum recall while restricting the total predicted positives to fewer than 50% of the data (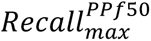).

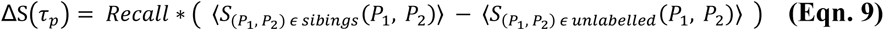

1. The difference in similarity scores (ΔS) between orphan sibling and unlabeled pairs indicates the distance between their score distributions. That is the difference in scores of functionally similar and unlabeled, most frequently not functionally similar, pairs (**Eqn. 9**). We weighted ΔS by corresponding methods’ Recall values to penalize methods for failing to identify test set siblings. ΔS varies in range [-1,1], where a positive value indicates higher similarity scores for sibling vs. non-sibling pairs. Note that ΔS reflects the distance between siblings and unlabeled pairs in the linear space of similarity scores. As a result, ΔS is comparable only among methods with similar similarity-score distributions. Despite its limitations, however, ΔS serves as a useful threshold-independent performance measure. Here, we report *ΔS_max_*, the highest ΔS for each method over the range of prediction score thresholds (τ_p_).

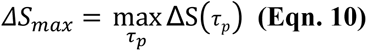
2. We also report the maximum Recall (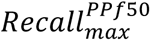) of each method over the thresholds (τ_p_ and τ_s_) while restricting the total predicted positives to fewer than 50% of the method-specific dataset. This measure reflects the best possible recall for each method, without encouraging trivial, i.e. “all pairs are siblings” positive overprediction.

We also compared the method prediction performances to two empirical random estimates: a random classifier and a random annotator. A random classifier samples the similarity scores (S) of protein pairs from a random uniform distribution. The random annotator is the result of shuffling sibling/unlabeled labels for all protein pairs in the test sets of each method in our assessment. Each simulation was repeated 100 times.

Further, the statistical significance of the performance differences between the tools in terms of ΔS, F_1max_, PR-AUC and ROC-AUC were assessed through the Wilcoxon rank-sum test and Student’s t-test using SciPy^80^.

## RESULTS AND DISCUSSION

### Assessing test set and evaluation metrics

We first aimed to evaluate our proposed strategy for assessing protein function annotation methods (Figure 2). Our protein structure alignment (Foldseek^54^) *plus* shared-function prediction (SNN^53^; **Figure S1**) based approach (SNN+TM; Methods) captures functional identity of a protein pair, labelling them *siblings*. Structural similarity is often used as a proxy of functional similarity^81, 82^. By filtering SNN predictions to structurally similar proteins, we assured high precision of our method.

To evaluate SNN+TM predictions, we annotated as siblings 1,927 (0.11%) of 1,745,646 protein pairs among 1,869 enzymes experimentally labelled with an Enzyme Commission^32^ (EC) numbers. For this set, all protein structures were predicted using ESMFold^39^ and only high-confidence structures were retained (**Methods**).

Note that our definition of siblings does not specifically reflect protein sequence similarity. The TM score component of our method is derived from alignment of the predicted protein structures. The SNN similarity score is predicted by a model that was trained to identify gene pairs encoding proteins from same *fusion* function clusters^53^. Proteins in different fusion clusters are often sequence similar (homologous), while proteins within the same cluster can be sequence dissimilar. To support our view that SNN predictions are not homology-driven, we observe that for our set of 1.7M enzyme pairs there was no correlation (−0.036) between the SNN score and sequence identity (**Figure S2**).

For this annotated enzyme set, we computed the performance of the “ground truth predictor” (**Table S1**), by under-sampling unlabeled pairs to generate siblings-to-unlabeled ratio of 1-to-1 over 100 iterations. In building our SNN+TM method, we selected the TM and SNN score thresholds (≥0.7 and ≥0.98, respectively) to attain ∼90% precision in capturing functionally similar proteins of our original labelled dataset^53^. That is, these cutoffs ensured that most protein pairs labelled as siblings are correctly labelled, but only a small fraction of all siblings is identified. Thus, for any set of proteins, our approach generates a dataset of positive (sibling) vs. unlabeled protein pairs, where the latter may contain siblings but, trivially, significantly fewer of them than non-siblings. Here, only 4% (69,042) of the unlabeled set (1,743,719 pairs total) were same EC pair proteins. At the same time, 88% (1,693) of the positive (sibling) set (of 1,927 pairs) had same EC numbers.

For this type of test sets (positive vs. unlabeled) the recall of assessed methods, i.e. their ability to identify positives/siblings, is justifiably the primary choice of performance measure. However, to avoid overestimating performance of methods that tend to overpredict positives, we needed to factor in the total number of positives labelled – a measure well captured by precision, i.e. the number of siblings predicted positive vs. all positive predictions. We note, however, that reported method precision values may be underestimated for some methods that correctly identify siblings from the unlabeled set. We thus use precision here only to illustrate the total number of positive predictions necessary for each method to recall known siblings.

A random classifier (**Methods**) can be expected to attain both recall and precision of 50% for a balanced (1:1) dataset. On the other hand, as expected, the recall and precision of our “ground truth predictor” were much higher – 88% and 96%, respectively **(Table S1 & S2).** Note that unlike with real function prediction methods that provide a confidence score with their prediction, for these experimental annotations we used binary, i.e. same function vs not, labels inferred from third digit EC number identity between protein pairs (**Methods**).

### Orphan siblings as a test dataset

From the MGnify^48^ collection of metagenomic data, we collected a set of 11,444 proteins having no homology (<30% identity) to any of the UniRef100 sequences (*orphans*) and paired them by expected SNN+TM functional identity (*orphan siblings*, Methods; **Figure 2**). As all supervised function prediction methods have been directly or indirectly trained on protein sequences found in UniProt, our approach eliminated any overlap between the training dataset of the prediction methods and our test dataset. Thus, our evaluation is an unbiased estimate of functional prediction method performance on any novel proteins.

Of the ∼65M possible protein pairs made from this set, only ∼309K attained a TM-scoreable structural alignment and 6,219 pairs (∼2%; **Methods**) were labelled functionally similar orphan siblings by our SNN+TM approach. This annotated dataset was used to assess the performance of protein function prediction tools.

Among the 309K protein pairs, 99.3% (307,434) shared <40% sequence identity, while 0.08% (240) were ≥80% identical (**Figure 2C**). Of the 6,219 orphan siblings, 5,576 (89.7%) had <40% identity and 95 (1.53%) were ≥80% identical, i.e. a slight enrichment for sequence similarity among orphan siblings as compared to the complete set of orphans. Despite this enrichment, however, siblings were largely composed of sequence dissimilar protein pairs.

### No one method is best for function annotation

We measured the ability of existing molecular functional annotation methods to assign identical functions to each protein in an orphan sibling pair (**Figure 3**, **S3 & Table 1; Methods)**. Note that methods differed in the number and kinds of proteins they could annotate, resulting in different test sets.

The top performers in this evaluation were not restricted to any one class of annotation; i.e. ECPred, GOPredSim, Pfam HMMER, and GhostKOALA attained similar performance using the ΔS, ΔS_max_, F1_max,_ etc. metrics (**Table 1 & Figure 3**). Curiously, except for GOPredSim, which uses ProtT5 embeddings^37^, the deep-learning models built to predict GO terms did not top the list. While NetGO attained solid performance, though still lower than the best method, other deep-learning methods (DeepFRI, DeepGOPlus, GOProFormer, and ProteInfer) were significantly lower (Wilcoxon rank-sum test, all p-values < 1E^-8^). Note that GOProFormer was developed on yeast proteins^60^ and using it to predict microbial protein function may have been beyond its scope of work.

Comparing performance measures derived at fixed thresholds (τ_p_^opt^ and τs^opt^) may not accurately depict the landscape of method performance. For a more stable measure, we computed the maximum of performance metrics (**Figure S3**) over a range of both prediction score and similarity score thresholds (τ_p_ and τ_s_). We also computed *ΔS_max,_* i.e. the linear distance between the distributions of sibling (positives) and non-sibling (negatives) similarity scores – a measure independent of any thresholds (**Figure 3A**). Using these metrics, ECPred, GOPredSim, Pfam HMMER, and GhostKOALA retained their position as top performers; additionally, Mantis Pfam predictions attained similar performance to the top scorers. Interestingly, performance of all methods was somewhat closer to the respective estimates of random than to the expected performance of the ideal ground truth predictor, highlighting the scope for improvement in function annotation.

We also computed the maximum recall (*Recall*_max_^*PPf*50^) of all methods across thresholds (τ_p_ and τ_s_) while limiting the number of predicted positives to ≤50% of the dataset (**Figure 3B**). This constraint restricts the inclusion of low confidence predictions and trivial overprediction of positives. GOPredSim and NetGO outperformed all the other annotation methods in this analysis. However, due to the differences in the number of proteins that could be annotated by each method, NetGO did so for a much larger number of sibling pairs. We thus note that though GOPredSim consistently performed well in all our analyses, the fraction of siblings it identified is significantly lower than other methods (**Figure 3B**).

Language models ESM, LookingGlass, and ProtT5 had similar, *Recall*_*max*_^*PPf*50^ as other methods (**Figure 3B & S4**). However, the respective performances of the method-specific random annotators (**Figure 3B**) for these three models were even higher. We are limited to speculating whether this results from the multi-dimensional and non-discrete nature of embeddings that encapsulate multiple protein characteristics, including structure similarity, homology, sequence length, etc., instead of function alone.

### Protein embedding distances are not directly informative of functional similarity

We further computed similarity between protein pairs based on cosine and Euclidean distances between embedding vectors of all 309K protein pairs (**Eqn. 4, 5**). Surprisingly, none of these distances captured the functional similarity between protein pairs well. Using our measures of performance (ΔS and *Recall_max_^PPf^*^50^), none of the language models did better than random. We note that when evaluation uses other metrics, specifically those relying on prediction precision, ESM-2, ProtT5, and LookingGlass embeddings achieve performance similar to some of the better predictors (**Figure S4**). However, as mentioned earlier, precision is not a reliable metric for our type of positive/unlabeled test data. Further, note that the generic choice of architecture did not drastically differentiate performance — among the top three embeddings, ESM-2 and ProtT5 are transformer-based protein language models^37, 39, 83^ whereas LookingGlass is a bi-directional LSTM model trained on short DNA reads^55^. The other four embeddings were SeqVec (LSTM) and Word2Vec (neural network) inspired by Natural Language Processing^72, 73^, Bepler is also a bi-directional LSTM trained on amino acid sequences^70^ and CPCProt is a convolution encoder trained through contrastive learning to identify subsequent fragments of protein against random protein fragments^71^.

We also note that given that ESM embeddings could predict protein structures, we expected these to capture functional similarity as defined, in part, by structural alignments. However, interpretation of language models is complicated and the extraction of average representation from multiple layers or a representation of any particular layer is bound to lead to loss of information^84, 85^. Our results thus highlight limitations of cross-domain application of unsupervised deep learning models without extensive analysis and fine-tuning.

### Function prediction strongly linked to protein structure

Our definition of siblings is dictated by structural similarity (TM alignment) and functional similarity (SNN). By using a high SNN cut-off (0.98), we have negotiated significant reduction in false positives at a loss of true positives. In other words, our approach to identifying functional siblings, while being very accurate (87.8% precision), is known to miss many protein pairs annotated to be of the same function (2.4% recall). While our structural similarity cutoff of TM score≥0.7 is an accepted value^86^ and the SNN cut-off of score≥0.98 was confirmed by our earlier experiments^53^, we aimed to explore method behaviour across the complete range of protein similarities. We thus computed the prediction performance of methods by varying the TM (**Figure 4**) and SNN score (**Figures S5**) cut-offs and, thus, redefining the protein pairs considered siblings. Note that in evaluating structural similarity we focused on pairs identified by Foldseek as possibly alignable (**Methods**), i.e. 309K protein pairs of 65M possibilities, but for these we varied the TM-score in the [0,1] range. The SNN method was trained to recognize pairs as functionally similar above the 0.5 cutoff, so we explored the SNN scores in the [0.5,1] range.

The top-performing methods (ECPred, GOPredSim, HMMER and GhostKOALA) showed consistently better performance than other methods across different TM and SNN score cut-offs. As expected, we observed a steep rise in performance of all methods at TM-score≥0.7 confirming that structural similarity above that threshold plays a significant role in functional similarity. At the same time, restricting the SNN score cutoff to 0.98 increased the performance (AUC under Precision-Recall curve and F1-score) of ECPred, ProteInfer, ProtCNN and DeepGOPlus by at least 20% (**Figure S6**) but reduced the performance of GhostKOALA. SNN captures functional similarity independent of either sequence or structural similarity^53^. We thus expect that tightening the SNN threshold reduces the performance of methods with strong dependency on sequence similarity, such as GhostKOALA.

Similarly, we repeated our assessment by varying the test dataset. We restricted our analysis to a subset of the test dataset consists of 4,376 protein pairs (including 1,700 siblings) made up of 3,506 proteins with no more than 150 residues to strictly restrict to single domain proteins (**Table S4**, **Figure S6 & S7**). Overall, we observed a slight in increase in performance of all GO and EC prediction methods except for GOPredSlim. Performance of GOPredSim significantly increased as measured by all the performance metrics, especially ΔS which nearly tripled. In contrast, methods predicting PFAM domain and orthologs should a slight decrease in performance. To our surprise, performance of GhostKOALA reduced drastically as observed in the previous observation.

### What do the best performing methods capture?

To answer this question, we evaluated contributions of known functionally relevant factors, i.e. sequence and structure, to method functional annotations.

We first clustered our orphan proteins, by sequence identity at different cut-offs between 0.9 to 0.4 using CD-HIT^52^ and explored their shared functionality (**Table 2, S5, S6 & Figure 5**). We found that less than a tenth of a percent of protein pairs in our orphan dataset set (240 pairs) were highly sequence-similar (80-90% seq.id), while the vast majority were not; i.e. more than 99% of protein pairs shared less than 40% sequence identity. Among all method predictions we observed significant enrichment of sequence similar protein pairs (≥40% seq. id.) and a depletion of dissimilar pairs. Note that this observation is unrelated to the putative correctness of their functional annotations.

**Fig. 5:**
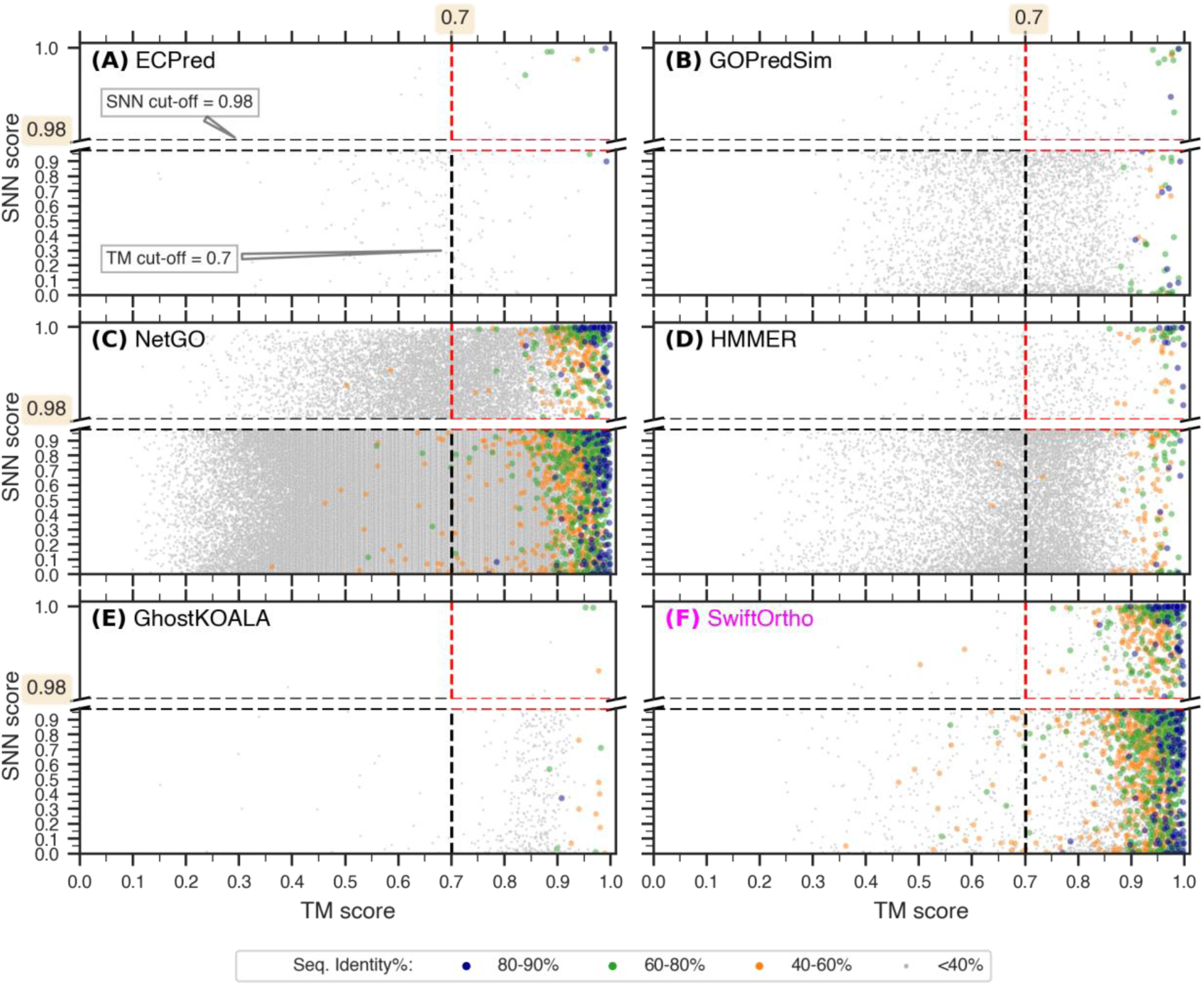
Different annotations capture different functional spaces. Orphan siblings predicted by the top-performing methods **(A-E)** and SwiftOrtho **(F)** occupy different spaces in the TM score (x-axis) vs SNN score (y-axis) space depending on the type of annotation (EC, GO, Pfam and orthologs). Each dot in the plot represents a protein-pair and is colored based on the sequence identity. The dashed lines indicate the TM and SNN score cut-offs (0.7 and 0.98) chosen for the defining the SNN+TM test set of labelled siblings (red dashed lines).

**Table 2:**
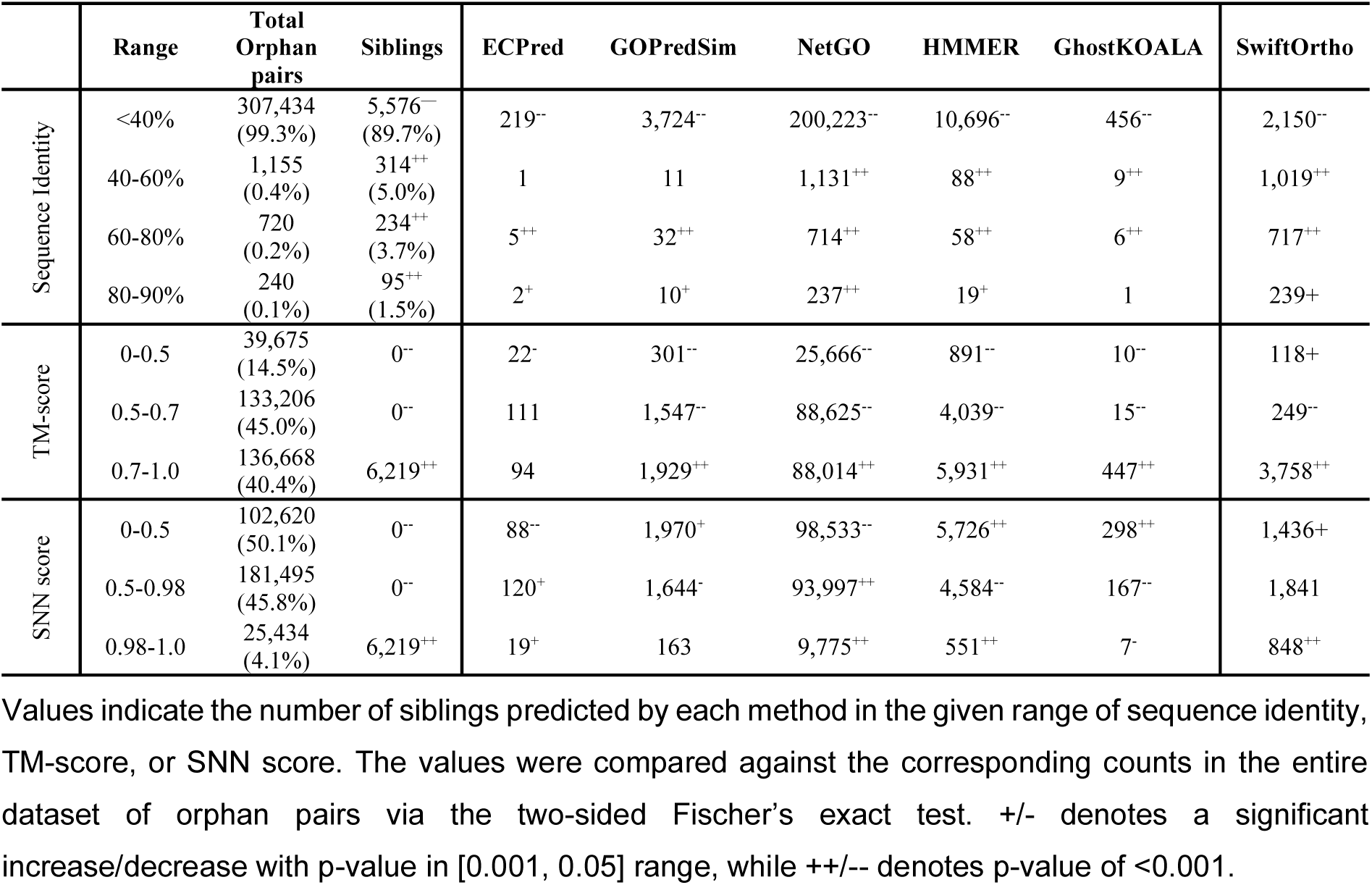
Characteristics of protein pairs predicted as siblings.

We further observed that the enrichment of GOPredSim and ECPred predictions was limited only to pairs of higher similarity (≥60%). GhostKOALA’s predictions, on the other hand, were not significantly enriched in highly sequence similar pairs (80-90%). Note that the small number of these highly sequence similar pairs complicates inference. That is, of the 240 such pairs, GOPredSim and ECPred identified ten and two, respectively – a small, but significant number – while GhostKOALA, which is built to annotate proteins using ortholog information, identified only one. We expect that the latter result is due to our test set being made up of orphan proteins, i.e. those without homologs in the predictor’s reference database. These observations suggest that, as expected, sequence information is important in driving function annotation by all methods, but GOPredSim and ECPred are more reliant than others on high sequence similarity.

The enrichment in the number of siblings predicted by each method within sequence-similar bins did not correlate with higher function prediction accuracy across these bins. For example, ECPred precision was worse for high similarity pairs than for lower ones, i.e. the opposite of the enrichment trend (**Table 2 and S6**). On the other hand, GOPredSim precision was similar for all sequence identity bins. HMMER which showed significant enrichment in the 40-80% sequence identity bin, had the highest precision of 63% in the 80-90% identity space. To summarize, while the predictions of the methods in this study are biased towards identifying sequence similar proteins as siblings, the accuracy of such predictions in terms of the functional similarity of the two does not agree with this assessment.

We also explored this homology-based functional annotations using SwiftOrtho^74^ – a method that identifies orthologous pairs in a given set of proteins based on sequence similarity^75, 76^. As expected, SwiftOrtho correctly identified 239 of 240 of the highly-sequence similar (80-90% sequence identity) pairs; it’s predictions were also enriched in pairs of sequence similar proteins at all levels of similarity ≥40% (**Table 2 & S5**). These results highlight the success achievable by homology-based methods in the presence of the relevant reference sets and further emphasize their deficiency in the absence of such reference. We note, however, that distinguishing orthologs from paralogs is hard^87, 88^ and even harder without the taxonomic and/or genomic context. In fact, SwiftOrtho also (putatively) incorrectly labelled 144 protein pairs as siblings.

We further analysed our data by binning predicted siblings based on structural (TM-score) and putatively functional similarities (SNN score). Predicted siblings from all methods, except ECPred were enriched in high structurally similar pairs (TM-score =[0.7,1.0]) vs. the low similarity range (TM-score =[0,0.5], **Table 2**). In other words, while most methods capture functional similarity driven by structural similarity, ECpred identified similar enzymatic activity in remotely structurally similar protein pairs as well. Note that ECPred predictions were significantly enriched in the high SNN score space ([0.98,1]). This is not unexpected given that functional convergence is more probable than sequence or even structural convergence^89, 90^ and ECPred relies on (sub)sequence and physiochemical feature similarity to predict EC numbers^68^.

Different patterns yet were observed in bins reflecting moderate levels of structural similarity (0.5-0.7) and sequence identity (40-60%). GhostKOALA showed a five-fold enrichment in protein pairs with moderate levels of sequence identity and a 14-fold depletion in protein pairs with moderate levels of structural similarity; a similar trend was observed for HMMER. Note that protein pairs with a TM-score over 0.5 are highly likely to share fold-level similarity^86^ – a feature that can be expected to reflect function, but does not appear to be useful to the methods reported here.

To summarize, sequence similarity is widely recognized as a key determinant in assessing functional similarity, particularly due to many functional evolution events resulting from gene duplication^91^. However, we hypothesize that existing protein sequence and domain recognition-based methods are biased towards capturing sequence similarity over functional signatures, thus failing to capture analogous evolution, reflect on function diverged between sequence-similar homologs, and identify conserved function among highly diverged siblings^24, 92, 93^.

Even considering the incomplete and erroneous functional annotations, everything we currently know about specific proteins and their functions is only a minor fraction of the entire protein universe^94^. However, annotating new proteins based on available data seems to be an inherently flawed proposition. Of the 53 million high-quality predicted ESM structures of microbial proteins extracted from MGnify only 54K (0.1%) had less than 30% sequence identity with UniRef. In turn, the overlap between all of UniProtKB and MGnify is estimated to be less than 1%^48^. That is, quality structure predictions, even with the aid of protein Language Models (pLMs), are limited to known protein families restrained by homology. For orphan proteins, this could explain the lacking performance when using embeddings (**Figure 3, S3 & S4)**. Nevertheless, GoPredSim which leverages function-transfer based on embedding similarity through k-nearest neighbors, is one of our four top performers, underscoring the potential of adapting current deep learning techniques to identify functional similarity among proteins.

### Summarizing the findings

With the growing stockpile of sequences, development of accurate functional annotation tools is more essential than ever. A major limitation to the assessment is the lack of large and diverse “ground truth” annotations. In this review, we assess some of the top protein annotation tools on a set of “orphan” proteins. To assess the quality of annotations, we translate the challenge of function prediction into a task of identifying functionally similar protein pairs in the dataset. Careful evaluation across a range of metrics reveals that even the performance of the top methods (ECPred, GOPredSim, HMMER and GhostKOALA) on this data is lacklustre. We note that even though the methods considered herein use different annotation vocabularies, our approach of deriving protein similarity scores enables their comparison.

In this review, we also explored the definition of protein functional similarity in terms of sequence and structure. Machine learning-based models such as ECPred, NetGO and GOPredSim capture more than sequence similarity from input sequences, unlike the more sequence homology-based algorithms. However, functional similarity is not only a function of sequence or structural similarity. The robustness of protein conformations paves way for diverse or similar sequences fold into diverse or similar structures to carry out the same or different functions as the environment dictates.

Another key observation from our work is that there is a lot of room for improvement in training deep learning models for protein functional annotation. Neither specifically trained methods, nor the cross-domain application of protein embeddings to identify functionally similar pairs showed promising results, highlighting the need for fine-tuning and analysis. While a huge advance has been made in protein structure determination in recent years, similar improvement in function annotation is limited by the lack of ground-truth annotation. However, alternate approaches to evaluation, such as the one we put forward, could pave the way for better models.

## Data availability

All data used in this study are listed in the main text or deposited in a permanent online data repository. The dataset of orphan proteins and the function similarity scores are available at <u>10.6084/m9.figshare.c.6737127</u>. The code used to compute siblings is available openly at https://bitbucket.org/bromberglab/siblings-detector/.

## Acknowledgements

We thank Iddo Friedberg (Iowa State University) and Predrag Radivojac (Northeastern University) for their thoughtful discussions on the topic of function. We are grateful to all the developers of methods presented herein for sharing their models and data and for responding to our questions. This work was supported by NAI Grant Number: 80NSSC18M0093. Y.B. was also supported by the NSF (National Science Foundation) CAREER award 1553289.

## Declaration of interests

The authors declare no competing interests.

## Notes

https://doi.org/10.6084/m9.figshare.c.6737127

### Competing Interest Statement

The authors have declared no competing interest.

### Summary of Updates

We have clarified the mechanism of siblings identification and the evaluation of functional annotation based on siblings in detail.

