## Supplementary material for "Functional profiling of the sequence stockpile: a review and assessment of *in silico* prediction tools"

**Prabakaran R[1,2*], Bromberg Y[1,2*]**

*1* Department of Biology, Emory University, Atlanta, GA 30322, USA
*2* Department of Computer Science, Emory University, Atlanta, GA 30322, USA

### Supplementary Online Material

#### Supplementary Text

##### The definition of Orphan siblings

In this work, a protein pair is defined as functionally similar or sibling pair based on the SNN+TM method[1], i.e. structural similarity as defined by TM score ≥0.7 and the SNN functional similarity score ≥0.98. The dataset of proteins used in our study is subset of proteins in the MGnify database of metagenome-assembled genomes[2] with <30% sequence identity with UniRef proteins[3]. In other words, our dataset consists of 11,444 orphan proteins with no homology to any annotated proteins in existing databases. The following pseudo-code explains the protocol used to annotate protein pairs as siblings. The respective code has been deposited at https://bitbucket.org/bromberglab/siblings-detector/.

```
Steps to identify siblings
-----------------------------------------------------------
Step 1: Compute TM score
a) Fetch ESM predicted structure for each protein[a]
b) Filter for structures with pLDDT and pTM > 0.90
c) Align structures with each other using FoldSeek[5]
d) Compute TM scores
-----------------------------------------------------------
Step 2: Compute SNN score
a) Fetch gene sequence for each protein
b) Calculate SNN score for each gene pair
-----------------------------------------------------------
Step 3: Annotated Orphan siblings
For each protein pair with TM score and SNN score
    If TM score ≥ 0.7 and SNN score ≥ 0.98 Then
        Annotate as "sibling"
    Else
        Annotate as "unlabelled"/"non-sibling"
    EndIf
```

[a]ESM structures can be fetched from ESM atlas or predicted using ESMFold[4].

### Tables

#### Table S1: Evaluation of experimental enzyme annotation

| Ratio of siblings to non-siblings | # of siblings | # of non-siblings | ΔS | Precision | Recall | Specificity | F1 score | Accuracy | Balanced Accuracy | MCC |
|---|---|---|---|---|---|---|---|---|---|---|
| 49:1 | 1927 | 39 | 0.739 | 0.999 | 0.879 | 0.963 | 0.935 | 0.880 | 0.921 | 0.340 |
| 9:1 | 1927 | 214 | 0.736 | 0.995 | 0.879 | 0.959 | 0.933 | 0.887 | 0.919 | 0.622 |
| 4:1 | 1927 | 481 | 0.736 | 0.989 | 0.879 | 0.960 | 0.930 | 0.895 | 0.919 | 0.739 |
| **1:1** | **1927** | **1927** | **0.737** | **0.957** | **0.879** | **0.960** | **0.916** | **0.919** | **0.919** | **0.842** |
| 1:4 | 1927 | 7708 | 0.737 | 0.848 | 0.879 | 0.961 | 0.863 | 0.944 | 0.920 | 0.828 |
| 1:9 | 1927 | 17343 | 0.737 | 0.711 | 0.879 | 0.960 | 0.786 | 0.952 | 0.919 | 0.765 |
| 1:49 | 1927 | 94423 | 0.737 | 0.312 | 0.879 | 0.960 | 0.460 | 0.959 | 0.919 | 0.509 |
| 1:99 | 1927 | 190773 | 0.737 | 0.183 | 0.879 | 0.960 | 0.303 | 0.960 | 0.919 | 0.390 |
| 1:499 | 1927 | 961573 | 0.737 | 0.043 | 0.879 | 0.960 | 0.081 | 0.960 | 0.919 | 0.188 |

We used experimental annotation as the ground truth function terms and evaluated these annotations for enzymes in UniProt database through our proposed SNN+TM approach. Values listed in the above table reflect the average performance of the ground truth annotations over 100 iterations.

#### Table S2: Evaluation of a random baseline classifier

| Ratio of siblings to non-siblings | # of siblings | # of non-siblings | ΔS | *ROC AUC* | *PR AUC* | Precision | Recall | Specificity | F1 score | *F1 max* | Accuracy | Balanced Accuracy | MCC |
|---|---|---|---|---|---|---|---|---|---|---|---|---|---|
| 49:1 | 1927 | 39 | 0.000 | 0.500 | 0.980 | 0.980 | 0.500 | 0.499 | 0.662 | 0.990 | 0.500 | 0.500 | 0.000 |
| 9:1 | 1927 | 214 | 0.000 | 0.501 | 0.900 | 0.901 | 0.502 | 0.502 | 0.644 | 0.947 | 0.502 | 0.502 | 0.002 |
| 4:1 | 1927 | 481 | 0.001 | 0.501 | 0.801 | 0.800 | 0.501 | 0.499 | 0.616 | 0.889 | 0.500 | 0.500 | 0.000 |
| **1:1** | **1927** | **1927** | **0.000** | **0.501** | **0.501** | **0.501** | **0.500** | **0.501** | **0.500** | **0.667** | **0.501** | **0.501** | **0.002** |
| 1:4 | 1927 | 7708 | -0.001 | 0.499 | 0.200 | 0.199 | 0.499 | 0.500 | 0.285 | 0.334 | 0.499 | 0.499 | -0.001 |
| 1:9 | 1927 | 17343 | 0.000 | 0.500 | 0.100 | 0.100 | 0.500 | 0.500 | 0.167 | 0.182 | 0.500 | 0.500 | 0.000 |
| 1:49 | 1927 | 94423 | 0.000 | 0.500 | 0.020 | 0.020 | 0.500 | 0.500 | 0.038 | 0.040 | 0.500 | 0.500 | 0.000 |
| 1:99 | 1927 | 190773 | 0.000 | 0.500 | 0.010 | 0.010 | 0.500 | 0.500 | 0.020 | 0.020 | 0.500 | 0.500 | 0.000 |
| 1:499 | 1927 | 961573 | 0.000 | 0.500 | 0.002 | 0.002 | 0.500 | 0.500 | 0.004 | 0.004 | 0.500 | 0.500 | 0.000 |

Values listed in the above table reflect the average performance of a random baseline classifier over 100 iterations (See **Table S1** for details).

Table S3: **List of methods**

| S. No | Method | Reference | Source | Type |
|---|---|---|---|---|
| 1 | DeepFRI | [6] | https://beta.deepfri.flatironinstitute.org/ | Webserver |
| 2 | ECPred | [7] | https://ecpred.kansil.org/ | Standalone |
| 3 | GhostKOALA | [8] | https://www.kegg.jp/ghostkoala/ | Webserver |
| 4 | GOPredSim | [9] | https://github.com/Rostlab/goPredSim | Python package |
| 5 | HMMER | [10] | http://hmmer.org/ | Standalone |
| 6 | Mantis | [11] | https://github.com/PedroMTQ/mantis | Standalone |
| 7 | NetGO | [12] | https://issubmission.sjtu.edu.cn/ng2/ | WebServer |
| 8 | ProtCNN | [13] | https://github.com/google-research/google-research/tree/master/using_dl_to_annotate_protein_universe | Python package |
| 9 | ProteInfer | [14] | https://github.com/google-research/proteinfer | Python package |
| 10 | Bepler | [15] | https://github.com/sacdallago/bio_embeddings | Python package |
| 11 | CPCProtEmbedder | [16] | https://github.com/sacdallago/bio_embeddings | Python package |
| 12 | ESM | [4] | https://github.com/sacdallago/bio_embeddings | Python package |
| 13 | LookingGlass | [17] | https://github.com/ahoarfrost/LookingGlass | Python package |
| 14 | ProtT5 | [18] | https://github.com/sacdallago/bio_embeddings | Python package |
| 15 | SeqVec | [19] | https://github.com/sacdallago/bio_embeddings | Python package |
| 16 | Word2Vec | [20] | https://github.com/sacdallago/bio_embeddings | Python package |

Table S4: **Evaluation on a dataset of single domain proteins**

| Tool | Term | ΔS | $F1_{max}$ | PR AUC | ROC AUC | Precision | Recall |
|---|---|---|---|---|---|---|---|
| ECPred | EC1 | **0.21** | **0.69** | **0.75** | **0.61** | **0.56** | **0.93** |
| ECPred | EC3 | **0.15** | **0.68** | **0.74** | **0.58** | **0.55** | **0.92** |
| Mantis | EC1 | 0.06 | 0.63 | 0.68 | 0.56 | 0.53 | 0.81 |
| Mantis | EC3 | 0.13 | 0.62 | 0.64 | 0.60 | 0.59 | 0.68 |
| DeepFRI | GO | 0.06 | 0.46 | 0.62 | 0.64 | 0.64 | 0.39 |
| GOPredSim | GO | **0.39** | **0.75** | **0.79** | **0.75** | **0.67** | **0.88** |
| Mantis | GO | 0.12 | 0.61 | 0.62 | 0.60 | 0.57 | 0.70 |
| NetGO | GO | 0.03 | **0.53** | 0.55 | 0.55 | 0.52 | 0.67 |
| ProteInfer | GO | 0.02 | 0.33 | 0.44 | 0.55 | 0.62 | 0.24 |
| GhostKOALA | KO | **0.03** | **0.34** | **0.45** | **0.55** | **0.65** | **0.25** |
| Mantis | COG | 0.03 | 0.49 | 0.50 | 0.53 | 0.54 | 0.47 |
| HMMER | Pfam | **0.07** | **0.63** | **0.69** | **0.54** | **0.53** | **0.83** |
| Mantis | Pfam | 0.03 | 0.53 | 0.54 | 0.53 | 0.53 | 0.55 |
| ProtCNN | Pfam | 0.00 | 0.11 | 0.34 | 0.51 | 0.62 | 0.07 |

We evaluated the performance of the methods in a subset of our original test dataset, consists of only proteins with sequence length less than or equal to 150 residues. The dataset consists of 1700 siblings among 4376 protein pairs. Values listed are average over 100 iterations of under sampling (**Methods**). Performance of the selected, top-performing methods is identified in bold.

Table S5: **Characteristics of protein pairs predicted as siblings**

| | Range | Total Orphan pairs | Siblings | ECPred | GOPredSim | NetGO | HMMER | GhostKOALA | SwiftOrtho |
|---|---|---|---|---|---|---|---|---|---|
| Sequence Identity | <40% | 99.32 | 89.66 -- | 96.48 -- | 98.6-- | 98.97-- | 98.48-- | 96.61-- | 52.12-- |
| | 40-60% | 0.37 | 5.05++ | 0.44 | 0.29 | 0.56++ | 0.81++ | 1.91++ | 24.7++ |
| | 60-80% | 0.23 | 3.76++ | 2.2++ | 0.85++ | 0.35++ | 0.53++ | 1.27++ | 17.38++ |
| | 80-90% | 0.08 | 1.53++ | 0.88+ | 0.26+ | 0.12++ | 0.17+ | 0.21 | 5.79++ |
| TM-score | 0-0.5 | 14.55 | 0.0-- | 9.69- | 7.97-- | 12.69-- | 8.2-- | 2.12-- | 2.86-- |
| | 0.5-0.7 | 45.07 | 0.0-- | 48.9 | 40.96-- | 43.81-- | 37.19-- | 3.18-- | 6.04-- |
| | 0.7-1.0 | 40.38 | 100.0++ | 41.41 | 51.07++ | 43.51++ | 54.61++ | 94.7++ | 91.1++ |
| SNN score | 0-0.5 | 50.12 | 0.0-- | 38.77-- | 52.16+ | 48.71-- | 52.72++ | 63.14++ | 34.81-- |
| | 0.5-0.98 | 45.81 | 0.0-- | 52.86+ | 43.53- | 46.46++ | 42.21-- | 35.38-- | 44.63 |
| | 0.98-1.0 | 4.07 | 100.0++ | 8.37+ | 4.32 | 4.83++ | 5.07++ | 1.48- | 20.56++ |

Values listed are the percentages of predicted siblings in the given range of sequence identity, TM-score and SNN score. The values were compared against the corresponding counts in the entire dataset of orphan pairs via the two-sided Fischer's exact test. +/- denotes a significant increase/decrease with p-value in [0.001, 0.05] range, while ++/-- denotes p-value of < 0.001

Table S6: **Characteristics of protein pairs predicted as siblings**

| | Range | Total Orphan pairs | Siblings | ECPred | GOPredSim | NetGO | HMMER | GhostKOALA | SwiftOrtho |
|---|---|---|---|---|---|---|---|---|---|
| Sequence Identity | <40% | 307,434 | 5,576 | 9/219 | 79/3,724 | 4,592/200,223 | 347/10,696 | 3/456 | 242/2,150 |
| | 40-60% | 1,155 | 314 | 1/1 | 2/11 | 302/1,131 | 19/88 | 1/9 | 259/1,019 |
| | 60-80% | 720 | 234 | 4/5 | 7/32 | 233/714 | 20/58 | 2/6 | 232/717 |
| | 80-90% | 240 | 95 | 1/2 | 2/10 | 95/237 | 12/19 | 0/1 | 95/239 |
| Total | | 309,549 | 6,219 | 15/227 | 90/3,777 | 5,222/202,305 | 398/10,861 | 6/472 | 828/4,125 |

*Values listed are the counts of correctly predicted siblings of total identified by the method as siblings in the given range of sequence identity.

### Figures

#### SI Fig. 1

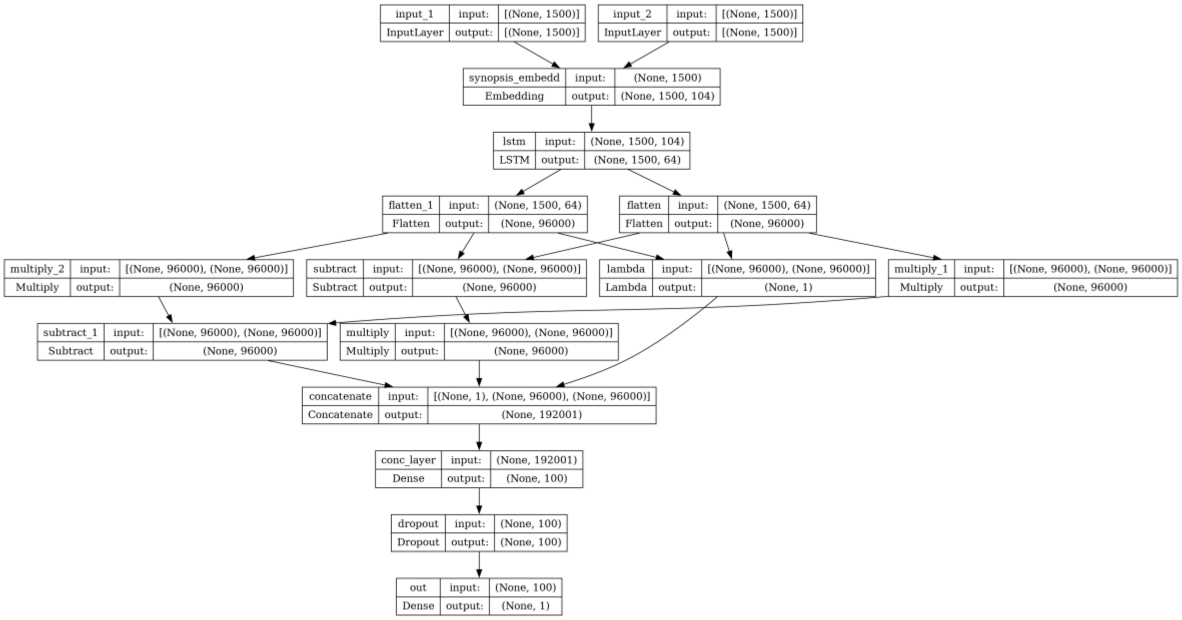


**SI Figure 1: Architecture of the Siamese Neural Network (SNN).** The flowchart describes the architecture of the Siamese Neural Network (SNN) model used in this study. The SNN architecture includes a) the pretrained embedding layer from LookingGlass[17], b) a LSTM layer and c) computation of distance between embeddings. The input consists of two protein-encoding genes, is split into two sequences of codons, and converted into embeddings through the LookingGlass embedding layer. Then the two sequences of embeddings are propagated through a layer of LSTM. Finally, the similarity between the two embeddings is computed as a combination of difference and cosine distance between embeddings and passed through a dense layer followed by sigma activation, which produces a similarity score.

#### SI Fig. 2

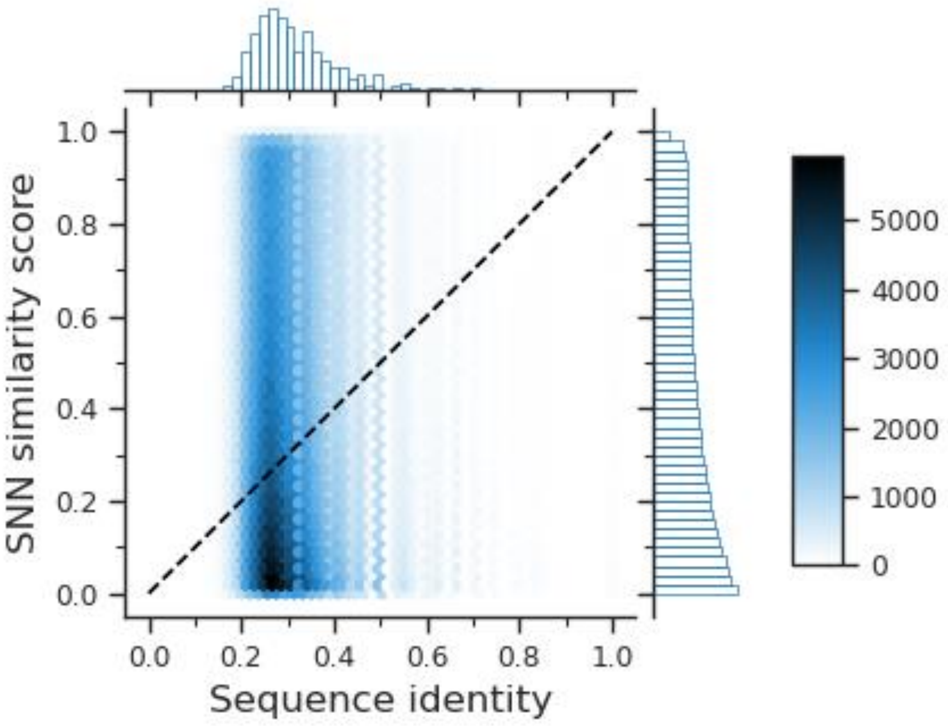


**SI Figure 2. Correlation between SNN score and Sequence identity.** The plot shows the distribution of sequence identities vs. SNN scores of 1,745,646 enzyme pairs used to validate our approach.

#### SI Fig. 3

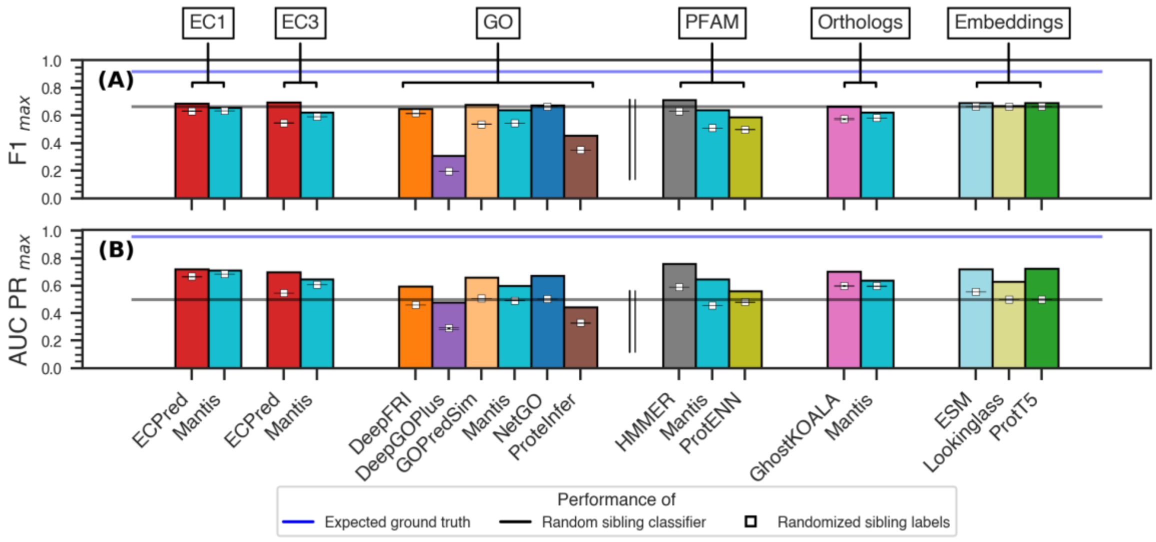


**SI Figure 3. Comparing method performance across embeddings.** The bar plots show the highest method (A) F1 scores, and (B) Area under the Precision-Recall curves for the prediction of orphan siblings (TM-score and SNN cutoffs of 0.7 and 0.98, respectively). The gray lines indicate the performance of the random classifier.

### SI Fig. 4

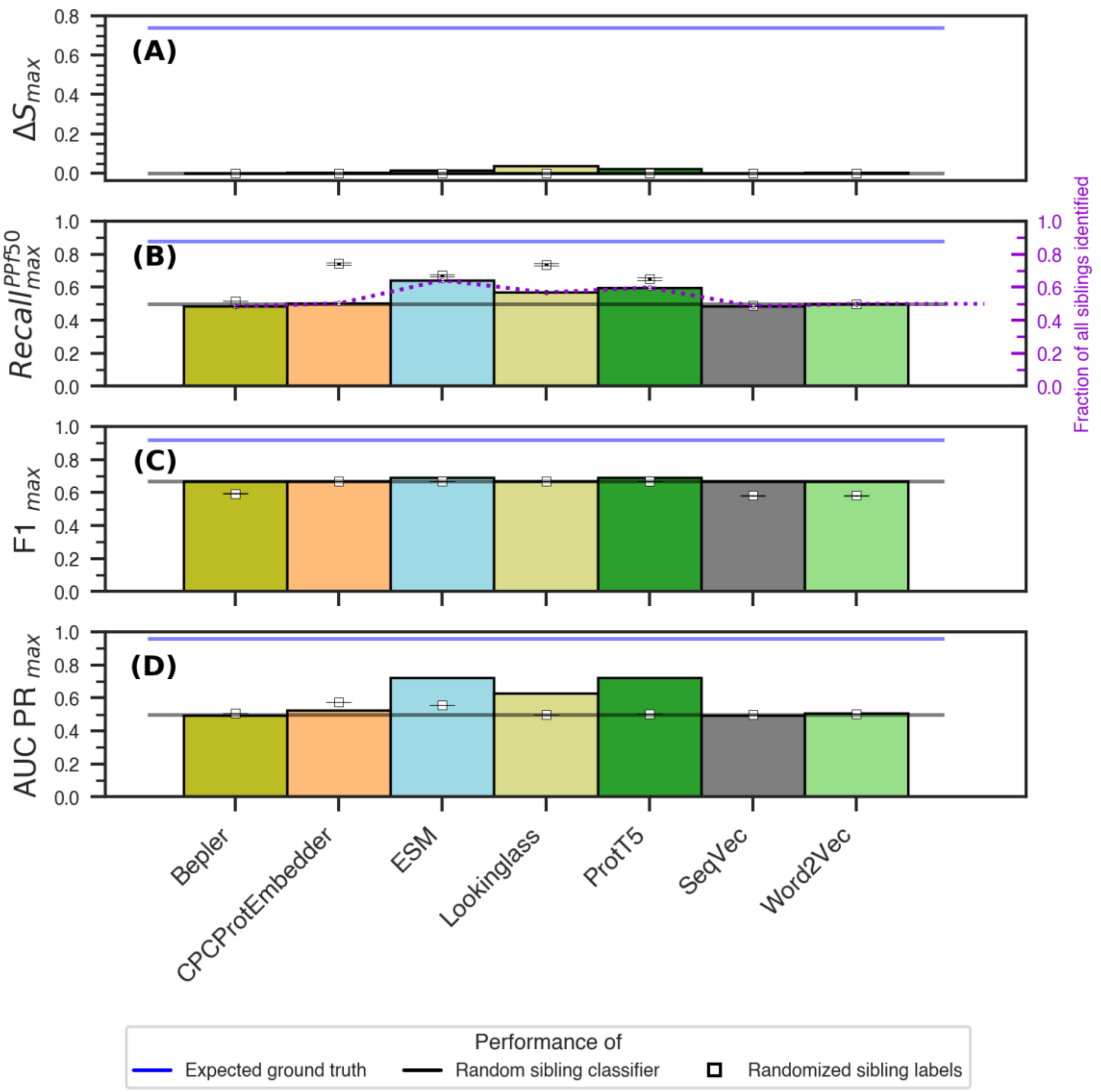


**SI Figure 4. Comparing method performance across embeddings.** The bar plots show the highest(max) method (A) ΔS, (B) $Recall^{PPf50}{}_{max}$, (C) F1 scores, and (D) Area under the Precision-Recall curves for the prediction of orphan siblings (TM-score and SNN cutoffs of 0.7 and 0.98, respectively). The gray lines indicate the performance of the random baseline classifier.

#### SI Fig. 5

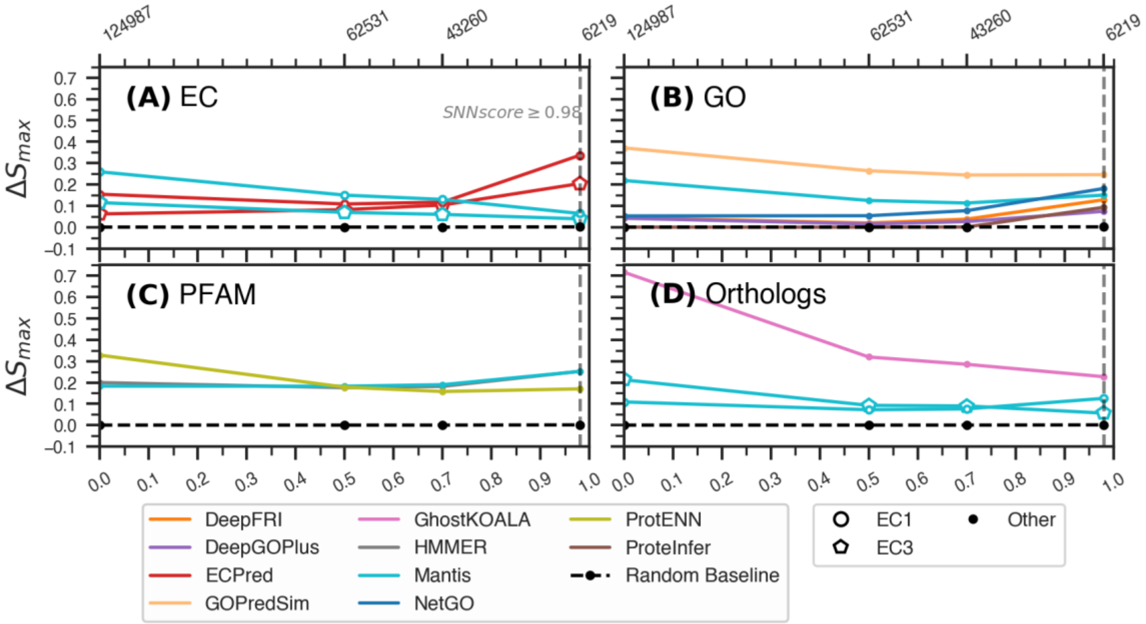


**SI Figure 5. Variations in ΔSimilarity score over SNN-score cutoffs.** The highest ΔS were computed at TM-score cutoff of 0.7 and at different SNN score cutoffs [0, 0.5, 0.7, 0.98] by varying the prediction score threshold. The average performance of 100 random baseline classifiers is plotted for comparison in each panel (black line).

**SI Fig. 6**

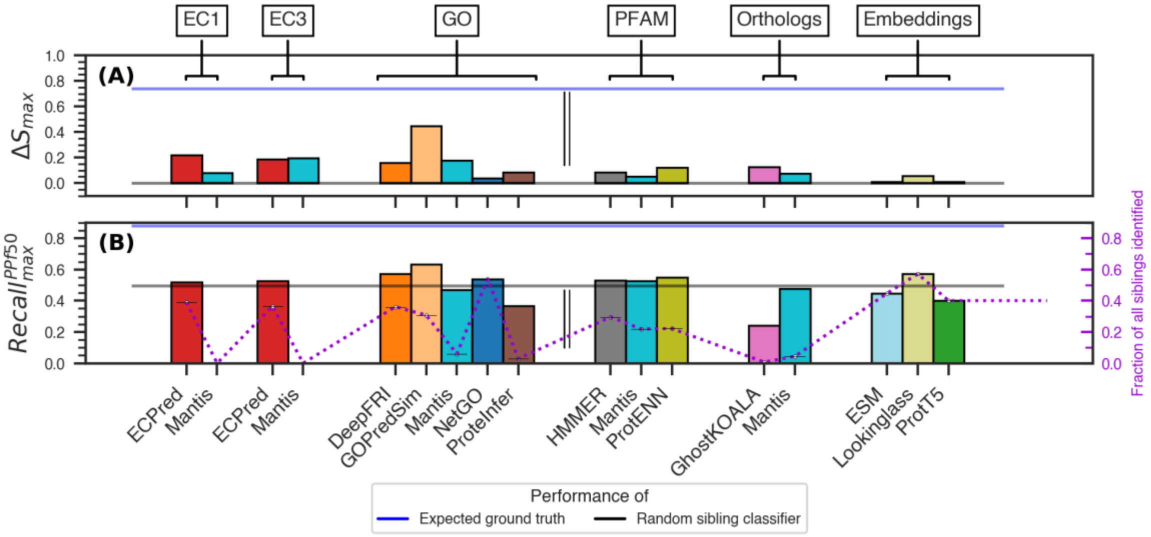


**SI Figure 6: Comparing method performance in each annotation category.** The bar plots show the highest (max) method **(A)** ∆S, and **(B)** $Recall^{PPf50}_{max}$ for the prediction of orphan siblings (TM-score and SNN cutoffs of 0.7 and 0.98, respectively), based on: Enzyme Commission numbers (EC1 – first digit, EC3 – up to third digit), Gene Ontology Molecular Function terms, Pfam domains, Ortholog groups and language model embeddings. The highest scores were selected for each method by varying the prediction score threshold. The double line separates explicit functional annotations (EC and GO) from Pfam and Ortholog definitions and embedding similarity. The gray lines indicate the average performance of 100 random baseline classifiers. The blue lines indicate the expected performance of the ground-truth annotation as observed in Table S1. Note DeepGOPlus was removed due to absence of sufficient protein pairs with predicted annotations.

#### SI Fig. 7

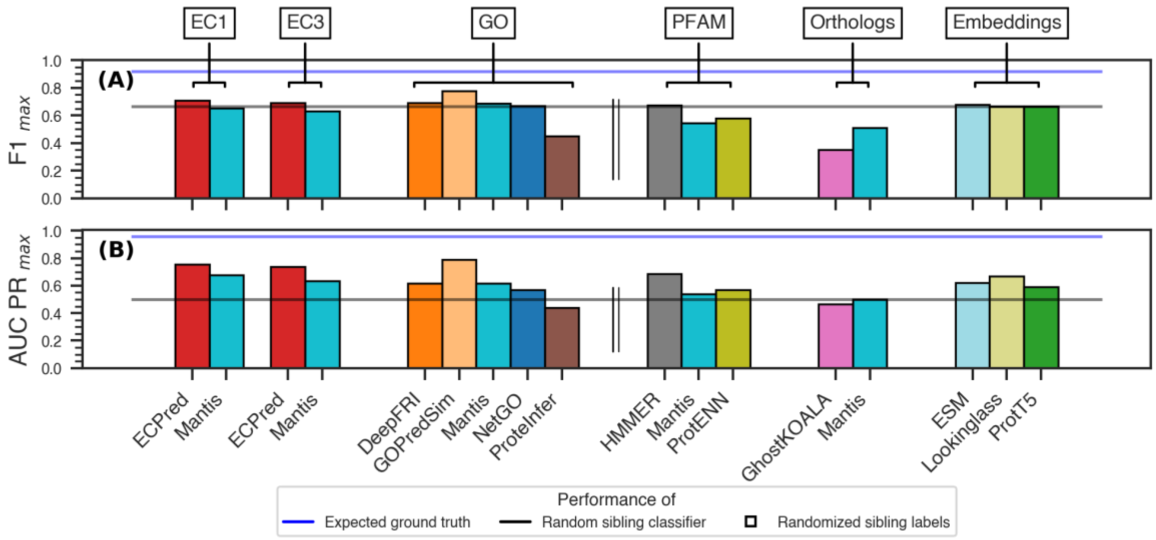


**SI Figure 7: Comparing method performance in each annotation category.** The bar plots show the highest (max) method **(A)** F1 scores, and **(C)** Area under the Precision-Recall curves for the prediction of orphan siblings (TM-score and SNN cutoffs of 0.7 and 0.98, respectively), based on: Enzyme Commission numbers (EC1 – first digit, EC3 – up to third digit), Gene Ontology Molecular Function terms, Pfam domains, Ortholog groups and language model embeddings. The highest scores were selected for each method by varying the prediction score threshold. The double line separates explicit functional annotations (EC and GO) from Pfam and Ortholog definitions and embedding similarity. The gray lines indicate the average performance of random baseline classifiers. The blue lines indicate the expected performance of the ground-truth annotation as observed in Table S1.
